# Associations between Infant-Mother Physiological Synchrony and 4- to 6-Month-Old Infants’ Emotion Regulation

**DOI:** 10.1101/2021.04.13.439498

**Authors:** Drew H. Abney, Elizabeth B. daSilva, Bennett I. Bertenthal

**Affiliations:** Department of Psychology, University of Georgia; Division of Science, Indiana University – Purdue University Columbus; Department of Psychological and Brain Sciences, Indiana University

**Keywords:** physiological synchrony, RSA, emotion regulation, biobehavioral development, Still Face Paradigm

## Abstract

In this study we assessed whether physiological synchrony between infants and mothers contributes to infants’ emotion regulation following a mild social stressor. Infants between 4- to 6-months of age and their mothers were tested in the Face-to-Face-Still-Face paradigm, and were assessed for behavioral and physiological self-regulation during and following the stressor. Physiological synchrony was calculated from a continuous measure of respiratory sinus arrhythmia (RSA) enabling us to cross-correlate the infants’ and mothers’ RSA responses. Without considering physiological synchrony, the evidence suggested that infants’ distress followed the prototypical pattern of increasing during the Still Face episode and then decreasing during the Reunion episode. Once physiological synchrony was added to the model, we observed that infants’ emotion regulation improved if mother-infant synchrony was positive, but not if it was negative. This result was qualified further by whether or not infants suppressed their RSA response during the Still Face episode. In sum, these findings highlight how individual differences in infants’ physiological responses contribute significantly to their self-regulation abilities.

---

Infants’ abilities to regulate their emotions undergo dramatic changes over their first year after birth. Early on, young infants demand a good deal of scaffolding or co-regulation from their parents, which they receive through face-to-face interactions, close physical contact, and vocal and affective turn-taking (e.g., Brazelton, Koslowski, & Main, 1975; Papousek, 1995). These behaviors are all examples of interpersonal synchrony, which refers to the “dynamic and reciprocal adaptation of the temporal structure of behaviors between interactive partners” (Leclère et al., 2014). As they mutually respond to each other’s signals, parent and child may coordinate not only their behavior and affective states, but also their biological rhythms and physiological responses (e.g. Feldman, 2007; Fogel, 1993). This coordination is deeply rooted within mammalian biology, and its success is foundational to the development of affiliative bonds and socio-emotional learning (Atzil, Hendler, & Feldman, 2014).

Currently, most studies focus on behavioral synchrony and its effect on both behavioral and physiological regulation. For example, infants of more synchronous dyads show greater affect regulation to a stressor, as measured by changes in affective behaviors and vagal tone (e.g., Moore & Calkins, 2004; Pratt, Singer, Kanat-Maymon, & Feldman, 2015). Mothers’ own physiological regulation of vagal activity also influences infants’ behavior and physiology (e.g. Leerkes et al., 2016; Moore et al., 2009). By contrast, most of what we know about how infant-mother physiological synchrony contributes to infants’ self-regulation is somewhat more speculative (Feldman, 2007, 2017). The goal of this study is to empirically test whether infant-mother physiological synchrony also contributes to infants’ physiological and behavioral regulation.

## Development of physiological synchrony

Humans are a social species from birth with the initial goal of regulating their physiological processes (or allostasis). Like other social species, human mothers engage in regulating not only their own physiological processes, but also in helping their infants adjust their internal states in order to grow and survive. Biobehavioral synchrony, including the regulation of temperature, immune function, and arousal, is one critical pathway for establishing allostasis that begins during gestation and continues after birth (Atzil, Gao, Fradkin, & Barrett, 2018). This form of synchrony involves the reciprocal coordination between behavioral, physiological, neural, and endocrine pathways both within and between individuals (Feldman, 2017), though these various forms of synchrony do not always align with each other (see Palumbo et al., 2017).

In this study, we focus on physiological synchrony involving vagal regulation of heart rate. When there are no challenging demands confronting the individual, the autonomic nervous system via the vagus nerve, attends to the internal viscera to maintain homeostasis and support growth and restoration (Porges, Doussard-Roosevelt, & Maiti, 1994). During sensory, cognitive and emotional challenges, the autonomic nervous system supports increased metabolic output by suppressing vagal output and increasing sympathetic excitation. According to Polyvagal Theory (Porges, 2003; 2007), different branches of the myelinated vagus nerve are responsible for regulating cardiac functioning as well as social engagement with the environment via eye contact, smiling, and vocalization. As the myelinated vagal fibers increase with development, visceral regulation improves and enables better and more independent behavioral regulation, especially with regard to social engagement behaviors (Porges & Furman, 2011). Prior to this developmental achievement, the mother assumes primary responsibility for regulating the physiological and behavioral state of the infant. Her ability to sense and respond to the physiological state of the infant is an important indicator of her sensitivity to maintaining the comfort, safety, and security of her infant (Feldman, 2007).

Given the goals of the current paper, it will be useful to provide a brief overview of the function and measurement of vagal tone. During rest or low demanding situations, vagal tone promotes homeostasis; the vagus acts as a brake to inhibit cardiac output. In contrast, when the organism is undergoing stress, the vagal brake is released (i.e. vagal suppression resulting in a decrease of vagal tone) and cardiac output is increased to facilitate mobilization behaviors (Bazhenova, Plonskaia, & Porges, 2001; Porges, Doussard-Roosevelt, Portales, & Greenspan, 1996). Vagal tone is typically measured as respiratory sinus arrhythmia (RSA) corresponding to heart rate variability within the frequency band of spontaneous breathing. Critically, higher vagal tone at baseline or rest is associated with greater adaptive functioning (e.g., social competence, emotion regulation, cognitive abilities) in response to environmental stimulation and challenges (Beauchaine, 2001; Blair & Peters, 2003; Porges et al., 1994). Both infants and young children with higher baseline RSA show a greater propensity for engaging in vagal suppression (Calkins, 1997; Huffman et al., 1998; Perry, Calkins, & Bell, 2016) and for regulating distress (Fox, 1989).

Human infants are not born with a fully-developed myelinated vagus. Rather, vagal development continues in the first few months postpartum (Fox, 1989; Porges & Furman, 2011), with the greatest increase in myelinated vagal fibers observed between 30-32 weeks gestational age and six months postpartum (Sachis, Armstrong, Becker, & Bryan, 1982). As a result of this strengthening of the vagal pathways, vagal tone steadily increases during the first few months after birth (Harper et al., 1977; Izard et al., 1991; Richards, 1989) and then stabilizes (Porter, Bryan, & Hsu, 1995). As the myelinated vagal fibers increase with development, visceral regulation improves and enables better behavioral and emotion regulation (Porges & Furman, 2011).

During early infancy, infants’ physiological regulation will often be tied to mothers’ co-regulation, which provides external support for their infants’ behavior. Repeated and ongoing synchronous interactions with caregivers along with an increase in myelinated vagal fibers contributes to the development of visceral regulation and enables the infant to express better behavioral regulation, which would include spontaneous social engagement behaviors (Porges & Furman, 2011). These periods of mutual engagement should be associated with increases in vagal tone for both infants and mothers to support homeostasis and maintenance of calm behavioral states (Bazhenova et al., 2001). Thus, it is hypothesized that greater infant-mother physiological synchrony should predict greater self-regulation for infants, as measured by increases in vagal tone and/or reduced distress or negative affect following a stressor.

## Infants’ responses to the Face-to-Face Still Face paradigm

We utilized a well-known social stressor, the Face-to-Face Still Face (FFSF) Paradigm (Tronick, Als, Adamson, Wise, & Brazelton, 1978) to determine how physiological synchrony contributes to infants’ behavioral and physiological regulation. The FFSF assesses infant-mother affective co-regulation by manipulating infant-mother interaction across three episodes. During *Social Play*, the dyad engages in face-to-face communication, followed by maternal *Still Face*, where the mother becomes completely non-responsive. Then, in *Reunion,* the mother re-engages with her infant and tries to repair the problems caused by the preceding episode.

The canonical response during FFSF is for infants to increase distress during maternal Still Face, and then decrease distress during maternal re-engagement (Mesman, Ijzendoorn, & Bakermans-Kranenburg, 2009). According to Polyvagal Theory (Porges, 2003, 2007), these behavioral changes should be accompanied by the suppression of vagal tone during the Still Face episode, enabling the infant to address the challenges created by the mother’s non-responsiveness. Typically, infants then increase vagal tone in Reunion once re-engagement with their mothers is re-established (Jones-Mason, Alkon, Coccia, & Bush, 2018; Moore et al., 2009).

Currently, there are inconsistencies in the literature regarding the extent to which behavioral and physiological responses co-occur. For example, Feldman, Magori-Cohen, Galili, Singer, & Louzoun (2011), report increased inter-beat-interval (IBI) synchrony of heart rate rhythms during moments of gaze, affect, and touch synchrony between infants and mothers.

However, infants have been found to show vagal suppression and subsequent recovery of vagal tone during and following maternal Still Face even while their negative affect persists upon maternal re-engagement (Moore & Calkins, 2004), thus demonstrating how physiological regulation can occur in the absence of behavioral regulation.

These contradictory findings may be attributable to the fact that behavioral and physiological responses were measured on different timescales or actually may be occurring at different timescales. Emotion regulation is a dynamic response that changes as a function of state and context (Cole, Martin, & Dennis, 2004), and therefore should be measured frequently. Yet, most studies rely on single time-averaged measures of distress or RSA over 30-120 seconds (Ham & Tronick, 2006; Pratt et al., 2015; Weinberg & Tronick, 1996; Conradt & Ablow, 2010). These aggregate measures can obscure dyadic adjustments occurring on faster timescales as both partners reciprocally respond to each other. In the current study, to better understand the dynamics of co-regulation, we will continuously measure social behaviors and RSA to assess the coherence between infants’ and mothers’ vagal activity during their face-to-face interaction and examine how this synchrony facilitates or hinders infants’ affect regulation.

## Positive vs. negative physiological synchrony

It is important to note that RSA synchrony, which is typically measured as a cross-correlation, can be positive or negative in direction (Abney, DaSilva, Lewis, & Bertenthal, under review). Positive synchrony refers to infants’ and mothers’ vagal activity changing in the same direction. Negative synchrony does not refer to the absence of synchrony, but rather infants’ and mothers’ vagal activity changing in opposite directions. Mothers and children have been observed to engage in positive physiological synchrony during face-to-face play and free play problem-solving tasks (Bornstein & Suess, 2000; Lunkenheimer et al., 2015; see Davis, West, Bilms, Morelen, & Suveg, 2018 for a recent review). Our primary goal was to evaluate whether positive compared to negative synchrony in the Social Play phase of the FFSF differentially predicts infants’ behavioral and physiological regulation. We were especially interested in the Reunion phase because of the unique regulatory demands it places on the dyad as they work to re-establish synchronous and reciprocal social interactions following the Still Face stressor (Cohn, 2003; Coppola, Aureli, Grazia, & Ponzetti, 2016).

This distinction between positive and negative synchrony was important for operationalizing our hypothesis: Infants experiencing positive physiological synchrony with their mothers during Social Play should show greater affect regulation post-stressor in the Reunion phase. Conversely, infants experiencing negative synchrony with their mothers during Social Play should have more difficulty regulating their negative affect during the Reunion phase (Weinberg & Tronick, 1996; Yoo & Reeb-Sutherland, 2013). In addition, we expect that dyads will differ in the degree of their physiological synchrony, just as they differ in the degree of their behavioral synchrony (Feldman & Eidelman, 2004; Granat, Gadassi, Gilboa-Schechtman, & Feldman, 2017); higher levels of physiological synchrony will be associated with greater reductions in negative affect as well as vagal reactivity (Provenzi et al., 2015; Woltering, Lishak, Elliott, Ferraro, & Granic, 2015).

## Individual differences in vagal suppression

Suppression of vagal tone during a challenge is an adaptive response that indicates the individual has physiologically adjusted to the environment and that there are available metabolic resources to efficiently engage in self-regulation of affect (Bazhenova et al., 2001; Porges et al., 1996). In general, infants and children who engage in vagal suppression, ‘suppressors’, show greater attentional control and soothability (Huffman et al., 1998) and fewer socially withdrawn and aggressive behaviors (Dale et al., 2006; Graziano & Derefinko, 2013; Porges, Doussard-Roosevelt, Portales, & Greenspan, 1996) compared to ‘non-suppressor’ infants whose vagal systems are less reactive. Although the overall trend is for infants to exhibit vagal suppression during challenges such as the maternal Still Face (Jones-Mason et al., 2018; Mesman et al., 2009), most studies report that only about 50% or fewer of the infants are suppressors who show the expected trend to decrease vagal tone during this period. Typically, these suppressor infants show greater affect regulation following an emotional challenge compared to non-suppressor infants (Bazhenova et al., 2001; Moore & Calkins, 2004; Provenzi et al., 2015).

Critically, these individual differences in the capacity for physiological regulation appear to interact with differences in infant-mother behavioral synchrony. For example, Provenzi et al. (2015) report that infants demonstrating a higher reparation rate from mismatched to matched behavioral states with their caregivers during Social Play revealed lower negative emotionality in Reunion, but this was only found for suppressor infants. One problem with interpreting these findings is that greater physiological regulation was confounded with suppressor infants exhibiting less distress during Still Face. As a consequence, it is difficult to determine if reduced distress during Reunion was a function of RSA suppression or their emotion state prior to the onset of Reunion. By contrast, Busuito et al. (2019) found the opposite effect: behavioral synchrony was inversely related to 6-month-old infants’ vagal tone, suggesting that infants with low vagal tone may have required greater support from their mothers in the form of behavioral synchrony. Likewise, Perry, Calkins, Nelson, Leerkes, & Marcovitch (2012) reported that 4-year-old children with lower vagal suppression were more influenced by maternal emotional socialization compared to children with higher vagal suppression who were capable of their own physiological regulation. Given the current state of the literature, additional studies are needed to better understand how infants’ physiological regulation interacts with infant-mother dyadic synchrony at both the physiological as well as at the behavioral level.

## Current Study

The primary goal of this study is to examine how synchrony of infant-mother vagal activity moderates infants’ behavioral and physiological regulation during the FFSF, with particular interest in the Reunion phase following the Still Face stressor. Given recent evidence of individual differences in the effects of behavioral synchrony on infants’ regulation of negative affect as a function of RSA suppression, we will also assess whether RSA suppression interacts with physiological synchrony. We hypothesize that infants who do not suppress vagal tone during the Still Face manipulation (i.e., non-suppressors) will require greater co-regulation as indexed by physiological synchrony with their mothers during the Social Play episode.

Physiological synchrony will be measured as both a continuous and categorical (positive vs. negative synchrony) variable, and we expect that greater evidence of synchrony will be associated with greater physiological and behavioral regulation for non-suppressor infants. By contrast, we anticipate that suppressor infants will engage in physiological and behavioral regulation regardless of their level of physiological synchrony with their mothers, because they are more capable of their own self-regulation and therefore depend less on maternal co-regulation.

Infants between 4- to 6-months of age were tested in order to capture the developmental period when individual differences in the capacity for behavioral and physiological regulation are emerging (Moore & Calkins, 2004; Porges et al., 1994; Provenzi et al., 2015). By including a dynamic measure of RSA and sampling behavioral measures continuously, we were able to analyze RSA and behavioral changes occurring within as well as between phases. These analyses were facilitated with the inclusion of growth curve models.

## Methods

### Participants

One hundred fourteen mothers and their 4- to 6-month-old infants (58 males) participated in the study. These participants were part of a larger longitudinal study on the development of self-regulation and were selected if electrocardiogram (ECG) data were available from both infant and mother. Families were part of a community sample recruited in a Midwestern college town. Overall, the sample reflects the local county demographics, which according to the most recent census is 83.4% White, 7.0% Asian, 3.6% Black, 3.5% Hispanic and 2.5% Multi-racial or other (US Census Bureau, 2018). Mothers were highly educated (i.e. college degree or higher) and ranged in age from 23- to 42-years-old (*M* = 31.36, *SD* = 4.22). All infants were born within 37-41 weeks gestational age. The Indiana University Institutional Review Board (IRB) approved all protocols for the study, “Infants’ social engagement and understanding” (#1310656881). Previous literature on analytic power using growth curve analyses suggests that sample sizes equal to or higher than 100 are appropriate (Curran, Obeidat, & Losardo, 2010).

### Procedure

Upon arriving at the lab, the family was greeted by a female experimenter who placed the infant in an infant seat to get acclimated before testing began. Mothers were seated in a chair facing the infant approximately 30 cm away. Two video cameras— one focused on the child and the other on the mother— were positioned unobtrusively in the corners of the room to record the session. A third camera (webcam) was used for the experimenter to monitor the dyad throughout the study.

The experimenter explained all procedures and obtained informed consent from the mother. Then, infant and mother were each fitted with three disposable Ag/AgCl ECG electrodes in a triangle pattern on their chests to measure their electrocardiographic responses continuously throughout the study. The electrode leads were hidden under the infant’s clothing and seat cover to minimize their attracting attention. The mother wore a wireless electrocardiogram (ECG) transmitter (Biopac Systems Inc., Goleta, CA) that did not restrict her movement. All recording was conducted using the Biopac MP150 system (Biopac Systems Inc., Goleta, CA). Following set-up, the experimenter sat behind a curtain for the duration of the study and monitored the dyad via webcam.

At the beginning of the study, the baby was placed in an infant seat and tethered around the waist to ensure that she could not climb out. The mother sat in a chair facing her infant.

During a 30-sec ECG baseline period, mothers completed a brief questionnaire; babies sat comfortably in their seat and visually explored the testing room. The FFSF paradigm consisted of three sequential phases that were verbally cued by the experimenter. In the **Social Play** phase, mothers were instructed to interact normally with their infants for two minutes. To minimize interference with heart rate recordings, mothers were told to not touch their infants (Feldman, Singer, & Zagoory, 2010). During the **Still Face** phase, mothers became unresponsive and posed a neutral expression staring at their infant’s forehead while keeping their face and body still for up to two minutes. Afterwards, mothers attempted to repair their relationship with their child and resume normal social interactions during a two-minute **Reunion** phase. If at any time infants became too distressed (i.e., hard crying for 10 seconds), the session was terminated early at the discretion of the mother or experimenter. In order to better measure the dynamics of the infant-mother interaction, each two-minute phase was divided into four 30-sec quartiles, for a total of 12 quartiles across the three FFSF phases.

### Data Coding

#### Observational codes

Infants’ and mothers’ facial expressions (smiling, distress, neutral, not scoreable), vocalizations (none, negative, non-negative, also speech for mothers) and gaze (mother, face/body, hands/objects, away and not scoreable) were coded using The Observer software (Noldus Information Technology Inc., Leesburg, VA). As described in Table 1, facial distress was characterized by tension between eyebrows, grimacing, the corners of mouth turned down and tension in lips (lip pursing). Negative vocalizations included fusses, whines, and cries; all other vocalizations including babbling, cooing, squealing, grunting, and laughing were scored as non-negative. Gaze aversion was operationalized as looking away from the social partner, including looking anywhere around the room, as well as at one’s own body. Codes within each category were mutually exclusive and exhaustive such that one code was always active per category.

**Table 1.** Behavioral coding definitions for vocalizations, facial expressions and gaze.

| Code | Description |
| --- | --- |
| **Facial Expressions** |  |
| Neutral/interest | Relaxed face, little to no tension; no emotion but engaged; no clear positive or negative emotion; also includes mouth movements such as lip smacking, chewing, blowing raspberries, tongue protrusion, kissy face, very specific movement/action, involved sneezes, coughs, yawns, etc. |
| Smiling | Corners of mouth pulled in and/or turned up, eyes squinted, cheeks raised, mouth open or closed |
| Distress | Tension between eyebrows, grimace, corners of mouth turned down, tension in lips |
| Not scoreable Face | Face obscured or not enough information to tell |
| **Vocalizations** |  |
| No vocalizations | No vocalizations (including pause in the mother's speech stream > 3 seconds); also includes sneezes/coughs |
| Non-negative | Babbling, cooing, squealing, grunting, laughing, gasping, etc. |
| Negative | Negative in valence; includes crying and whimpering (when coding baby) |
| Speech | Talking or singing (when coding mother) |
| **Gaze** |  |
| Face/Body | Looking at social partner's face or body |
| Hands/Objects | Looking at social partner's hands or nearby object (including infant seat) |
| Gaze aversion | Looking away from social partner, including looking down or at one's own body or gazing around the room |
| Gaze not scoreable | Eyes closed or obscured, or not enough information to tell |

#### Reliability

Reliability was calculated for 15% percent of all videos. In the Observer program, codes were evaluated for code matching and duration. Raters achieved the following percent agreement for infant behaviors: vocalizations (81%), facial expressions (86%), and gaze (76%). To control for chance agreement, kappa scores were also calculated. Raters achieved substantial agreement (Landis & Koch, 1977) across all categories: vocalizations (K= 0.64), facial expressions (K= 0.81) and gaze (K= 0.68).

#### Behavioral distress

A composite measure of distress was created based upon the amount of time infants exhibited negative facial expressions, negative vocalizations, and gaze aversion. In order to credit infants for expressing multiple negative behaviors per second, scores were computed as follows. If infants expressed only one behavior, they received a score of “1”; two behaviors resulted in a score of “2”, and three behaviors resulted in a score of “3”. Scores were summed within quartiles, which was then divided by the duration of the quartile (typically 30 sec), to compute a composite distress score. For example, the score for an infant crying during 15 sec of the quartile was 0.5 (15/30), whereas an infant crying for 15 sec and looking away for 30 sec was 1.5 (15 + 30) / 30). The maximum score an infant could receive is 3 (90/30) if they were displaying negative facial expressions, negative vocalizations, and gaze aversion for the entire duration of the analyzed time period. Thus, scores could range from 0 to 3, and were computed for each of the 12 quartiles across the FFSF; final scores were divided by 3 so that all values corresponded to a 0-1 scale. For infants who terminated the Still Face (n = 4) or Reunion phase (n = 10) early, missing quartiles were imputed using the previous quartile’s score.

### Physiological data processing

Both infants’ and mothers’ electrocardiograms were recorded at a sampling rate of 1 KHz. Trained research assistants detected and removed artifacts due to movement, ectopic beats, and periodic bradycardias. Editing was completed offline using CardioEdit software (Brain Body Center, University of Illinois at Chicago). Following the method developed by Porges (1985), the IBI time series was first time-sampled at regular intervals (5 Hz.), then a 51-point band-pass local cubic polynomial filter was used to estimate and remove the slow periodic and aperiodic components of the time-series. Finally, an FIR-type bandpass filter was applied that further isolated the variance in the IBI series to only the frequency range of spontaneous breathing for infants (0.3-1.3 Hz.) and for adults (0.12-1.0 Hz.). The higher range of 1.0 Hz for mothers’ respiration was used to account for the infrequent occurrence of faster breathing during talking or playing segments, so that the same filter could be used for all mothers in all conditions. The Porges & Bohrer (1990) technique for RSA magnitude estimation includes parsing this component signal into discrete epochs (lasting 10 to 120 sec), then calculating the natural log of the variance in each epoch. RSA is reported in units of ln(ms)^2^.

Unlike previous studies, we also modified this method to enable a more continuous exploration of the co-variance in the two estimates of RSA magnitude (mother and infant). At the magnitude estimation stage, a sliding window of 15 seconds was used to extract a continuous (updated every 200 ms) estimate of cardiac vagal tone for both participants. The estimated RSA value appeared at the beginning of the sliding window. Physiological synchrony was then estimated from the co-variation between these two estimates of time-varying cardiac vagal tone (see Abney et al. (under review) for more details concerning this procedure). All RSA time series data are available at the Open Science Framework: https://osf.io/udxn7/?view_only=65142acbb6b14def80b5f254d638b8eb.

## Results

We begin by describing how RSA synchrony was calculated and how dyads were categorized into positive vs. negative RSA synchrony groups. The division of the three FFSF phases into 12 quartiles enabled us to assess the dynamic changes in distress across the experiment. We conducted four sets of analyses. In the first set, we used growth curve modeling to test how infants’ behavioral distress changed as a function of two factors: Vagal Suppression (suppressors vs. non-suppressors), and RSA synchronization (positive vs. negative). (See below for details on classification criteria for both of these variables.) Based on the two hypotheses of this study, we were especially interested in the interaction between these two variables. In the second and third sets of analyses, we test whether the degree of physiological synchrony was associated with reductions in behavioral distress (second analysis) and physiological regulation (third analysis) following the Still Face phase. Similar to the first analysis, we also test whether these associations differ for suppressor and non-suppressor infants. In the last set of analyses, we test whether there were associations between behavioral and physiological regulation during the Reunion phase.

### Estimation of RSA synchrony and classification

RSA synchrony was based on infants’ and mothers’ RSA sampled at 5Hz during the Social Play phase. Both of these signals were detrended by computing the 1^st^-order derivative from the original signal and then they were submitted to a cross-correlation analysis where the lag-0 cross correlation was estimated. In the following analyses, we first treated synchrony as a categorical variable to determine how distress trajectories differed as a function of synchrony valence. Growth curve modeling was used to study distress trajectories over time as a function of synchrony. This analytic technique required that the synchrony variable was categorized. In the second set of analyses, we treated synchrony as a continuous variable using the lag-0 coefficient as the measure of synchrony.

In order to classify synchrony, we converted the cross-correlation values into a dichotomous variable (positive vs. negative synchrony depending on whether the lag-0 coefficients were above or below, *r*=0, respectively)^1^. As can be seen in Figure 1, the greatest difference between positive and negative synchronies occurred at lag-0, which is the reason we defined synchrony as the co-occurrence of the RSA values and not as one participant leading or lagging the other. The distribution of dyads categorized as either ‘positive’ (n=58 dyads) or ‘negative’ (n=56 dyads) was uniformly distributed.

**Figure 1.**
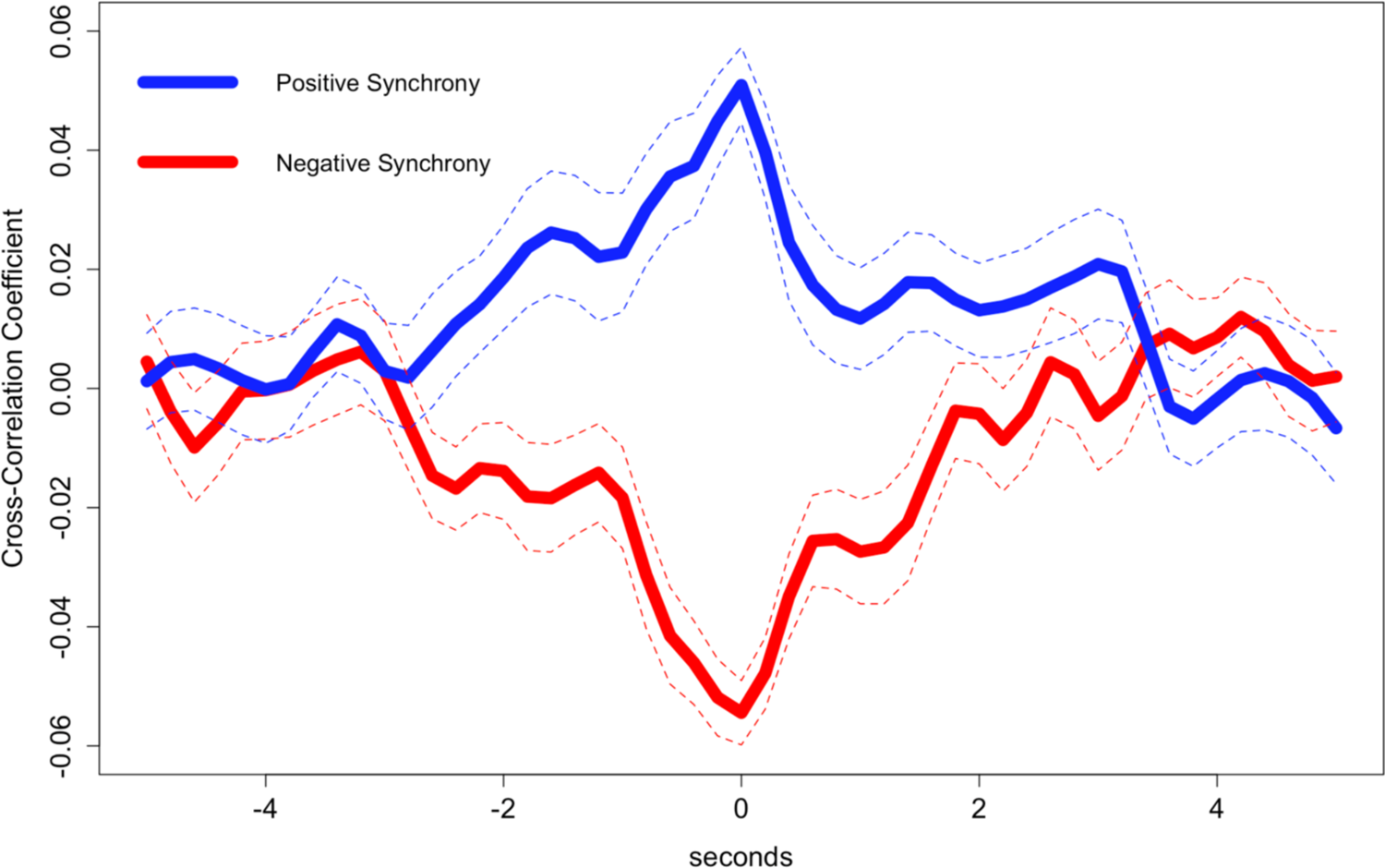
Average cross-correlation functions for dyads classified as positive (blue) or negative (red) synchrony. Dashed lines indicate +/-1 standard error of the mean.

### Testing Null Hypothesis of RSA Synchrony

It is important to determine whether or not our observations of synchrony go beyond patterns that could be explained by the natural frequency of RSA signals (a shuffled test) and also beyond what would be explained by randomly pairing dyad members (a surrogate test). We conducted a permutation test in order to ask: Do cross-correlation coefficients from empirical dyads come from a statistically similar distribution as empirical dyads after we shuffle phase-level information in the signal? To answer this question, for the positive synchrony group and negative synchrony group, separately, we randomly permuted infant and mother RSA signals and submitted the within-dyad shuffled signals to cross-correlation analysis and estimated the lag-0 cross-correlation coefficient. Non-parametric Kolmogorov-Smirnov (KS) tests compared the distributions of lag-0 cross-correlations from the empirical and shuffled dyads. The null hypothesis of the KS test is that the two distributions are sampled from a population with statistically similar distributions. The KS tests revealed that empirical and shuffled distributions were not sampled from the same population for the positive synchrony group (D = 0.60, *p* < .001) nor the negative synchrony group (D = 0.48, *p* < .001).

In the second set of analyses, we conducted a surrogate test in order to ask: Do cross-correlation coefficients from empirical dyads come from a statistically similar distribution as coefficients from random pairings of individuals into dyads? To answer this question, for the positive synchrony group and negative synchrony group, separately, we randomly paired an infant from one dyad with a mother from another dyad and estimated the lag-0 cross-correlation coefficient. Distributions of 1000 cross-correlation coefficients from surrogate pairs were estimated by iterating this process 1000 times for the positive and negative synchrony groups, separately. The KS tests revealed that empirical and surrogate distributions were not sampled from the same population for the positive synchrony group (D = 0.53, *p* < .001) nor the negative synchrony group (D = 0.51, *p* < .001). These results thus confirm that the empirical distribution of lag-0 cross correlations is not a function of chance.

### Analysis 1: Do changes in infants’ behavioral distress differ as a function of RSA suppression and RSA synchrony?

#### Description of Growth Curve Analyses

To test for differences in the trajectories of distress across the three phases, we used mixed-effects models and growth curve analyses as described in Mirman (2014). In growth curve analysis, the predictor variables are considered in terms of change over time (or ‘growth’). We specifically tested whether there were differences in the distress trajectories across the FFSF phases and whether trajectories were moderated by Vagal Suppression and/or RSA synchronization.

In the model, distress is the outcome variable and the linear and quadratic terms are used to predict distress. The canonical trajectory of distress in the FFSF paradigm is that it begins low in the Social Play phase, increases during the Still Face phase, and then decreases during the Reunion phase (Mesman et al., 2009). This distress trajectory is best explained by a negative quadratic polynomial (see Figure 2). As suggested by Mirman (2014), polynomial terms were generated orthogonally to allow for independent contributions of the linear and quadratic terms. This resulted in models corresponding to second-order polynomial regression models. We used the **lme4** library in R (Bates, Maechler, Bolker, & Walker, 2014) to construct linear mixed effects regression models. The models were maximally specified as long as the models converged. We used random intercepts (Dyad ID) and nested the n^th^-order polynomial terms (e.g., linear and quadratic).

**Figure 2.**
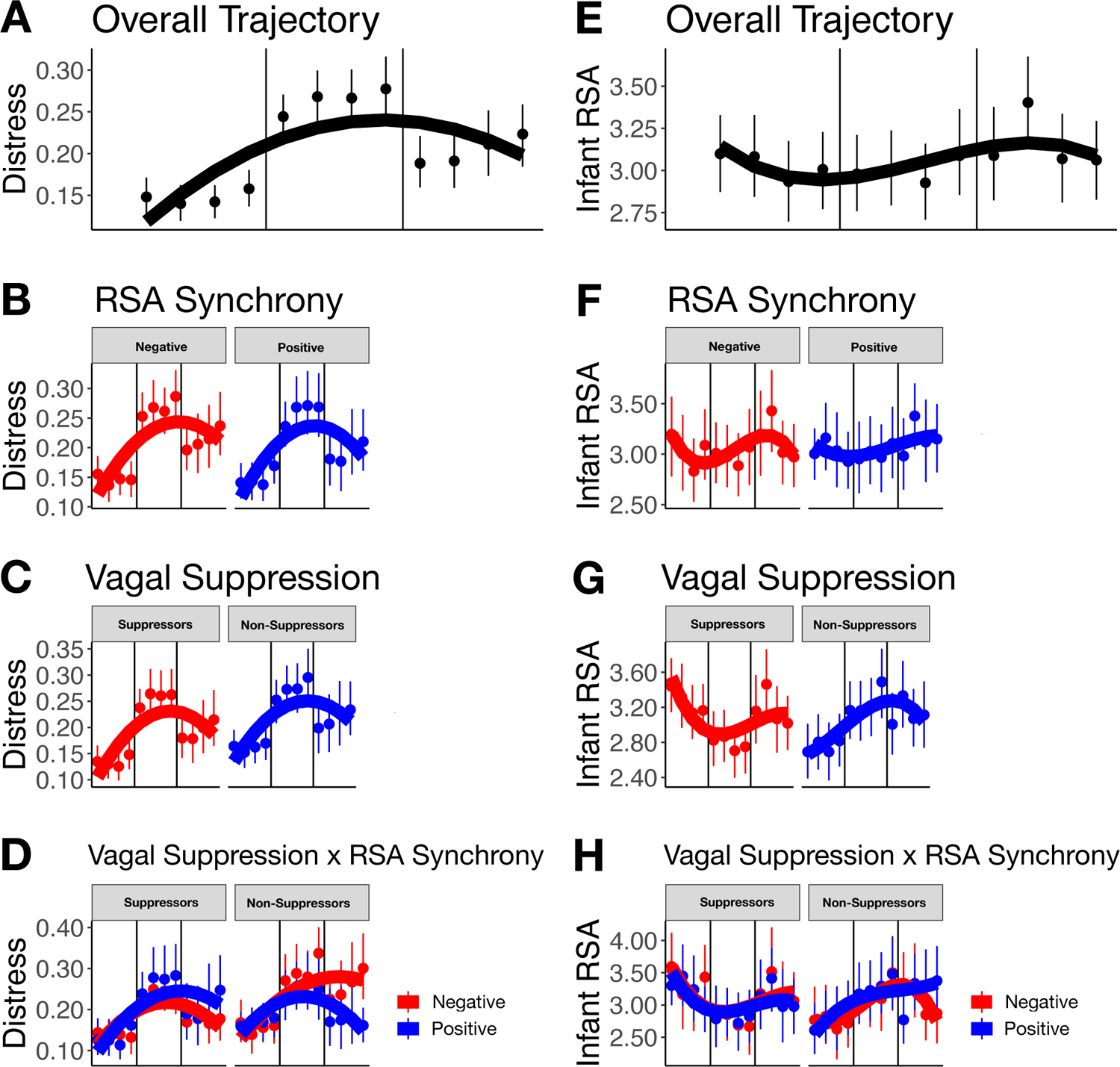
Left column. (A) The overall trajectory of infants’ distress across the three phases. The trajectory of infants’ distress across the three phases as a function of (B) RSA synchrony, (C) Vagal Suppression, and (D) the interaction between Vagal Suppression and RSA synchrony. Right column. (E) The overall trajectory of infants’ RSA across the three phases. The trajectory of infants’ RSA across the three phases as a function of (F) RSA synchrony, (G) Vagal Suppression, and (H) the interaction between Vagal Suppression and RSA synchrony. Solid lines reflect quadratic fit. Error bars indicate -/+95% confidence intervals.

The Vagal Suppression variable was created by classifying infants into either a positive or negative change score, depending on whether RSA increased or decreased from the Social Play phase to the Still Face phase, respectively (i.e. increases corresponded to no-suppression and decreases corresponded to suppression of RSA; Bazhenova et al., 2001; Pratt et al., 2015). Note, that this method is somewhat different from the method of comparing RSA during Still Face to RSA during the baseline (e.g. Moore & Calkins, 2004). The problem with using baseline RSA is that infants’ RSA is unlikely to remain stable in this period, because infants are just beginning to adjust to a new situation. Although infants’ RSA does not remain stable during the subsequent Social Play phase either, this measure is at least closer in time to RSA during the Still Face phase, and thus constitutes a more reliable measure of whether RSA increases or decreases during the Still Face phase (cf. Bazhenova et al., 2001). It should be noted that this problem is not unique to the FFSP, and there is considerable variability in trying to determine a reliable baseline for physiological studies (see Jones-Mason et al., 2018 for a longer discussion of these issues).

#### Descriptive statistics

The first set of analyses provide a summary of the measures used in the growth curve modeling. These measures include behavioral distress as the outcome measure and infants’ RSA synchrony and Vagal Suppression as the predictors. Table 2 summarizes the sample sizes, means, and standard deviations of these measures, and group comparisons. As can be seen, there were significant differences in RSA synchrony between the negative and positive synchrony groups and differences in mean RSA change score between the suppressor and non-suppressor groups. The differences in distress during the three phases of the Still Face were also significant and will be reported in the section on Overall Trajectory of Distress.

**Table 2.** Descriptive Statistics (sample size, means and standard deviations) and t-statistics for RSA synchrony, Vagal Suppression, and Behavioral Distress by Phase along with Baseline RSA as a function of RSA Synchrony or Vagal Suppression.

| Variables | Mean (SD) | t-statistic (Cohen's d) |
| --- | --- | --- |
| Baseline RSA |  |  |
| RSA Synchrony |  |  |
| Negative (n=56) | 3.00 (1.22) | 0.14 (0.33) |
| Positive (n=58) | 3.04 (1.16) |  |
| Vagal Suppression |  |  |
| Suppressors (n=52) | 3.11 (1.27) | 0.89 (0.17) |
| Non-Suppressors (n=62) | 2.91 (1.10) |  |
| RSA Synchrony |  |  |
| Negative (n=56) | -0.06 (0.04) | 11.73*** (2.19) |
| Positive (n=58) | 0.05 (0.05) |  |
| Vagal Suppression |  |  |
| Suppressors (n=52) | -0.50 (0.38) | 12.16*** (2.25) |
| Non-Suppressors (n=62) | 0.56 (0.54) |  |
| Distress by Phase |  |  |
| Social Play (n=114) | 0.15 (0.09) |  |
| Still Face (n=114) | 0.26 (0.15) |  |
| Reunion (n=114) | 0.20 (0.17) |  |

Note. \*\*\* $p < .001$

### Results of Growth Curve Analyses

In the following subsections, we report growth curve model results for the relationships between the functional form (linear and quadratic) of the trajectory of behavioral distress and Vagal Suppression and RSA Synchrony. We begin by testing whether infants’ distress follows the canonical trajectory for the Still Face paradigm (Figure 2A). Next we test whether RSA Synchrony moderates the distress trajectory. Following each of these analyses, we add Vagal Suppression to the model to evaluate whether the direction of RSA change during Still-Face interacts with the previous variable.

### Overall Trajectory of Distress

The overall trajectory of distress was tested with a growth curve model fit up to a cubic polynomial. As seen in Figure 2A, the canonical trajectory of distress was supported by the observation that the best fitting model was an orthogonalized polynomial model including a linear and quadratic term, (X^2^ = 67.59, *p* < .001). Including the cubic polynomial did not increase model fit, X^2^ = 0.03, *p* = .86. The negative quadratic model fit reflects the overall pattern of distress that increases from the Social Play phase to the Still Face phase and decreases from the Still Face phase to the Reunion phase. The results of the growth curve model were corroborated when submitting the phase-level estimates of distress to a one-way ANOVA, which showed a significant main effect of Phase, *F*(2, 226) = 35.51, *p* < .001, η_p_^2^ = .239. Post-hoc t-tests comparing distress estimates across the three phases indicated that distress increased from the Social Play phase to the Still Face phase (*t*[113] = −8.73, *p* < .001, Cohen’s *d* = .95), and then decreased from the Still Face phase to the Reunion phase, *t*(113) = 4.29, *p* < .001, Cohen’s *d* = .40. Finally, distress was significantly higher in the Reunion phase compared to the Social Play phase, (*t*[113] = −4.00, *p* < .001, Cohen’s *d* = .46, suggesting that distress attenuates gradually and requires a period of time before returning to the Social Play level.

### RSA Synchrony

There were no significant interactions between the functional forms of distress and RSA Synchrony, linear: β = −0.14, *p* = .55; quadratic: β = −0.09, *p* = .46 (see Figure 2B).

### RSA Suppression

There were no significant interactions between the functional forms of distress and RSA Suppression, linear: β = −0.02, *p* = .92; quadratic: β = 0.06, *p* = .65 (see Figure 2C).

### RSA Synchrony and RSA Suppression interaction

There was a significant three-way interaction between the linear trajectory of distress, RSA Suppression, and RSA synchrony, linear: β = −1.29, *p* < .001 (see Figure 2D). By contrast, the three-way interaction between the quadratic trajectory of distress, Vagal Suppression, and RSA Synchrony was not significant, β = −0.20, *p* = .43. As can be seen in Figure 2D, non-suppressor infants who were classified as engaging in positive RSA Synchrony demonstrated a decrease in distress during Reunion. By contrast, non-suppressor infants who were classified as engaging in negative RSA Synchrony demonstrated no reduction in distress during Reunion. In other words, non-suppressor infants were unable to regulate their distress during Reunion unless they had experienced positive RSA synchrony with their mothers during Social Play. By contrast, infants who had experienced negative RSA synchrony with their mothers during Social Play demonstrated a carry-over effect during Reunion whereby their distress remained high. The responses of the suppressor infants were similar to the non-suppressor infants through the Still Face phase, but there was no dissociation in their distress response during the Reunion phase.

### Infants’ RSA

We replicated the analyses conducted above but replaced changes in infants’ distress with changes in infants’ RSA. One noteworthy difference is that the best fitting polynomial was neither a linear nor quadratic trajectory, but instead a cubic trajectory (see Figure 2E), which suggests that, overall, there were two changes in direction in the trajectory of infants’ RSA throughout the three phases. There were significant interactions between the functional forms of infants’ RSA and Vagal Suppression, linear: β = 0.58, *p* < .001; quadratic: β = −0.68, *p* < .001. These interactions were driven by the Vagal Suppression variable: Infants’ RSA decreased for suppressors and increased for non-suppressors. All other models involving RSA trajectories, RSA synchrony, and Vagal Suppression revealed no significant differences (see Figures 2F-H). Although there were differences in RSA during Social Play (*t*(110) = 2.97, *p* = .001), it is unlikely that these differences are responsible for moderating the effects of RSA synchrony. If RSA level and RSA suppression were confounded, then the correlation between the two should have been significant for both suppressors and non-suppressors. The results revealed a significant correlation for the suppressors (*r*(60) = 0.43, *p* < .001), but not for the non-suppressors (*r*(50) = 0.06, *p* = .675).

### Analysis 2: Is the degree of physiological synchrony linearly associated with reductions in behavioral distress?

The results from the first set of analyses suggested that non-suppressor infants benefit from positive physiological synchrony with their mothers. In the first set of analyses, the synchrony variable was categorical, which precluded a clear test of an association between the strength of synchrony and changes in behavioral distress. To further explore this relationship, we ran zero-order correlations between the degree of RSA synchrony (as a continuous variable) in Social Play and the amount of change in distress that infants displayed from the Still Face to the Reunion phase. We computed difference scores for the four quartiles (30s each) of the Reunion phase by subtracting the mean distress scores for each quartile of the Reunion phase from the mean distress score during Still Face; more positive scores indicated greater reduction in distress. These difference scores were divided into two groups (suppressors and non-suppressors) and zero-order correlations were then calculated between each of the four difference scores and the RSA synchrony correlation coefficient calculated previously. To control for multiple correlations, we set our alpha to .05/8 correlations = .006. A simple power analysis suggested that with a sample size of *n*=52 (*n*_Suppressors_=52, *n*_Non-suppressors_=62), the power to detect a large effect size (*r*=0.5) given our adjusted alpha is .87.

Table 3 reports the zero-order correlations. Notably the only significant correlation between the RSA synchrony and reduction in distress was for non-suppressor infants during the final quartile of the Reunion phase, *r*(50) = .39, *p* = .004. This result suggests that the more strongly non-suppressor infants were positively synchronized with their mothers, the more they reduced their behavioral distress in the last 30 seconds of the Reunion phase.

**Table 3.** Zero-order correlations for RSA synchrony and changes in behavioral distress as a function of Vagal Suppression.

| Variables | Reunion: 1<sup>st</sup><br>Quartile | Reunion: 2<sup>nd</sup><br>Quartile | Reunion: 3<sup>rd</sup><br>Quartile | Reunion: 4<sup>th</sup><br>Quartile |
| --- | --- | --- | --- | --- |
| *Suppressors* | -0.01 | -0.13 | -0.17 | -0.12 |
| *Non-Suppressors* | -0.07 | 0.08 | 0.14 | 0.39* |

*Note.* \*adjusted alpha of $p < .006$

### Analysis 3: Is the degree of RSA synchrony associated with reductions in infants’ RSA?

Similar to the motivations of the second set of analyses, we explored how RSA synchrony is related to physiological regulation during Reunion for suppressor and non-suppressor infants. We computed difference scores for the four quartiles (30s) of the Reunion phase and mean RSA during the Still Face phase, and conducted zero-order correlations separately for suppressor and non-suppressor infants. To control for multiple correlations, we again set our alpha to .05/8 correlations = .006.

There was only one marginal result (i.e., not significant following correction for multiple tests) involving the correlation coefficient for RSA synchrony and changes in infants’ RSA during Reunion (RSA during Still Face – RSA during Reunion quartiles). As can be seen in Table 4, this result involved only the non-suppressor infants during the first quartile of the Reunion phase, *r*(50) = 0.31, *p* = .03. Given that this result does not meet the threshold for significance when corrected for multiple correlations, it is best not to try to interpret it.

**Table 4.**
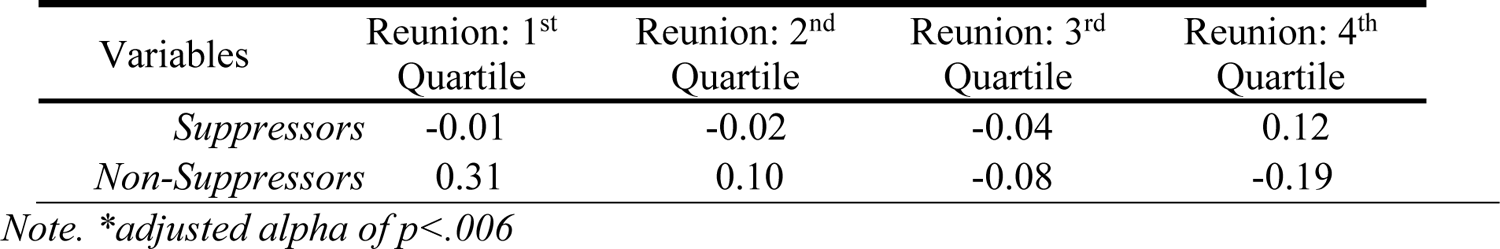
Zero-order correlations for RSA Synchrony and changes in infants’ RSA as a function of Vagal Suppression.

| Variables | Reunion: 1<sup>st</sup><br>Quartile | Reunion: 2<sup>nd</sup><br>Quartile | Reunion: 3<sup>rd</sup><br>Quartile | Reunion: 4<sup>th</sup><br>Quartile |
| --- | --- | --- | --- | --- |
| *Suppressors* | -0.01 | -0.02 | -0.04 | 0.12 |
| *Non-Suppressors* | 0.31 | 0.10 | -0.08 | -0.19 |

*Note.* \*adjusted alpha of $p < .006$

### Analysis 4: Is there a relationship between RSA and behavioral regulation?

In a final set of analyses, we tested whether changes in infants’ RSA were associated with changes in distress during the Reunion phase (see Table 5). To control for multiple correlations, we again set our alpha to .05/8 correlations = .006. As before, changes were quantified by subtracting the mean Still Face response from each of the four quartile responses during Reunion. Non-suppressor infants revealed a marginal association (i.e., not significant following correction) between changes in distress and RSA during the final quartile of the Reunion phase, *r*(50) = −0.27, *p* = .05.

**Table 5.** Zero-order correlations for changes in distress and infants’ RSA as a function of Vagal Suppression

| Variables | Reunion: 1<sup>st</sup><br>Quartile | Reunion: 2<sup>nd</sup><br>Quartile | Reunion: 3<sup>rd</sup><br>Quartile | Reunion: 4<sup>th</sup><br>Quartile |
| --- | --- | --- | --- | --- |
| *Suppressors* | -0.04 | 0.07 | -0.03 | -0.21 |
| *Non-Suppressors* | 0.07 | 0.10 | 0.09 | -0.27 |

*Note.* \*adjusted alpha of $p < .006$

## Discussion

The primary aim of this study was to understand the relation between physiological synchrony of vagal activity between infants and their mothers during Social Play and infants’ behavioral and physiological regulation following a stressor. Unlike most previous studies (e.g. Jones-Mason et al., 2018; Mesman et al., 2009), we used growth curve analyses to understand how trajectories of distress and RSA differed across the FFSF overall as a function of physiological synchrony and infants’ physiological regulation. When examining all the infants together, we observed the canonical ‘Still Face effect’ whereby infants’ distress increased during Still Face and then decreased in Reunion, i.e., a quadratic trajectory (Adamson & Frick, 2003; Jones-Mason et al., 2018; Mesman et al., 2009). Interestingly, the trajectory for RSA was not the inverse of distress, but rather it corresponded to a cubic trend whereby RSA tended to decrease through the beginning of the Still Face phase, followed by an increase that decreased toward the end of the Reunion phase. Previous Still Face studies quantifying RSA change more coarsely at the phase-level report a negative quadratic function decreasing in Still Face and increasing in Reunion (see Jones-Mason et al., 2018). By examining within-phase changes every 30 seconds, this study provides a more nuanced perspective regarding the temporal dynamics of RSA change across the FFSF phases.

Furthermore, our findings revealed that infants’ responses were moderated by their own physiological regulation. In our sample, 54% of infants were suppressors, and 46% were non-suppressors, which is comparable to what other studies report for 4- to 6-month-old infants (e.g. Bazhenova, Plonskaia, & Porges, 2001; Moore & Calkins, 2004; Provenzi et al., 2015).

Critically, we found that individual differences in physiological regulation – whether infants suppressed or did not suppress vagal tone during maternal Still Face – impacted the observed patterns of distress and RSA.

### Behavioral regulation of distress

Infants who did not suppress RSA during the Still Face episode and were classified as displaying negative synchrony with their mothers were unable to attenuate their distress during the Reunion phase. By contrast, this failure to regulate distress was not shared by infants classified as displaying positive synchrony with their mothers. Infants’ distress response continued to diminish throughout the Reunion episode. Apparently, mothers displaying positive synchrony with their infants were better able to co-regulate their infants’ behaviors, and compensate for their inability to regulate their own arousal precipitated by the Still Face manipulation. This result highlights how the previously documented ‘carry-over’ effect of distress into Reunion (e.g. Weinberg & Tronick, 1996) differs as a function of physiological synchrony, at least for non-suppressor infants.

Infants who suppressed RSA during Still Face demonstrated similar patterns of emotional regulation during Still Face and Reunion episodes regardless of whether they engaged in positive or negative synchrony with their mothers. In particular, infants in both groups demonstrated that negative affect diminished over time during the Reunion episode. Our interpretation for this result is that infants who suppressed RSA were better able to regulate independently the attentional, behavioral, and emotional demands associated with the Still Face stressor (Bazhenova et al., 2001; Degangi, Dipietro, Greenspan, & Porges, 1991; Gueron-Sela et al., 2017; Huffman et al., 1998; Moore & Calkins, 2004; Provenzi et al., 2015; Stifter & Corey, 2001). As such, they were less dependent on the external support provided by the mother via her synchronous activity.

Converging evidence for these findings was obtained with RSA synchrony measured not as a dichotomous variable, but as a continuous variable sampled at 5 Hz. This measure enabled us to quantify the strength of the coupling between infants’ and mothers’ vagal activity during their Social Play phase. The results revealed that there was a significant linear relation between the magnitude of RSA synchrony and the attenuation of distress during the Reunion episode, but this relation was only present for the non-suppressor infants. Moreover, this effect was not significant until the final quartile of the Reunion episode, suggesting that co-regulation of the infant’s negative affect emerged gradually following the Still Face episode. In sum, these results suggest that physiological synchrony, like behavioral synchrony, contributes to infants’ regulation of their affective state, but the effects are more likely to be observed with infants whose vagal system is less reactive.

One very important implication of these findings is that individual differences are critical for fully understanding the responses of infants following a social stressor. For example, the quality of mother-infant interaction is a crucial contributor to infants’ emotion regulation. In previous research, it was reported that dyads of non-suppressor infants are characterized by less than optimal mother-infant interaction, such as lower levels of dyadic synchrony (Moore & Calkins, 2004) and maternal sensitivity (Conradt & Ablow, 2010). Based on the current findings, it appears that we cannot predict the quality of mother-infant interactions for non-suppressor infants without taking into account physiological synchrony as well.

### Physiological regulation of distress

As we previously mentioned, past studies investigating changes in RSA during the FFSF paradigm report a trajectory that appears to be the inverse of the distress trajectory: RSA decreases from Social Play to Still Face, and then increases from Still Face to Reunion. By sampling RSA four times as frequently as done in previous studies, we observed that the best fitting polynomial corresponded to a cubic as opposed to quadratic trajectory. The main finding from this analysis was that the trajectories differed as a function of vagal suppression, which was expected given that RSA trajectories were, by definition, different during the Still Face episode.

It is also noteworthy that the main findings associated with changes in negative affect during Reunion were not replicated when the dependent measure was RSA. In particular, the RSA trajectory was not moderated by physiological synchrony nor by vagal suppression.

Moreover, RSA synchrony measured as a continuous variable did not predict either increases or decreases in RSA during the Reunion episode.

### Relation between behavioral and physiological regulation

It is generally assumed that changes in both physiological reactivity and negative affect are correlates of emotion regulation during the FFSF paradigm (e.g. Field, 1994; Haley & Stansbury, 2003; Mesman et al., 2009), but the empirical results tend to reveal significant inconsistencies. For example, Weinberg & Tronick, (1996) reported that during the Reunion episode, heart rate and vagal tone returned to levels observed during Social Play, suggesting that infants’ affective responses should also return to the same levels observed during Social Play.

Yet, the results were not consistent with this prediction. Infants were generally positive and showed little negative affect during Social Play. During Reunion, however, infants displayed a mixed pattern of both positive and negative affect, which was not expected given their physiological responses. According to Beauchaine (2001), the relation between physiological and behavioral regulation is complicated because RSA is associated with both negative and positive expressions of emotion. RSA appears to mark the capacity for active engagement of infants with the environment, but this engagement can be associated with either approach or avoidance.

This inconsistency between physiological and behavioral results was also observed in our study. Infants’ changes in RSA and negative affect were not correlated in any of the four quartiles during Reunion regardless of whether or not they were suppressors. One interpretation for these findings is that expressed negative affect and autonomic measures are not tightly coupled. According to Gunnar, Mangelsdorf, Larson, & Hertsgaard (1989) infants’ affective displays are specifically related to different interactive events, but their physiological reactions do not show the same level of specificity. It is also important to acknowledge that infants’ behaviors are multiply determined by complex interactions involving biological and psychological systems that are continuing to develop. The differences between suppressor and non-suppressor infants could very likely be attributable to different rates of development of the vagal pathways. Likewise, the differences among infants in their regulation of negative affect could be in part attributable to their social and cognitive development, which manages their expectations regarding maternal responsiveness (Bigelow & Power, 2014; Bigelow & Walden, 2009). Also, the quality of the affective communication between infants and mothers contributes to infants learning how to regulate their emotions (Tronick, 1989). All of these factors develop according to their own timetables and are largely determined by the individual experiences of each child and their mother. Thus, we do not expect these and other factors develop synchronously, which provides one compelling reason for why infants’ physiological and behavioral responses are not highly correlated.

One additional finding that may seem somewhat more puzzling concerns why physiological synchrony was more likely to predict behavioral than physiological regulation during the Reunion episode. The reason may be more related to measurement issues than anything else. Although the correlation between the magnitude of RSA synchrony during Social Play and RSA change scores was not significant, the lack of significance may have been attributable to the compression of RSA scores during Reunion relative to the range of distress scores, which was 10 times as great. As a consequence, the likelihood of observing a significant correlation was decreased.

### Limitations and Future Directions

The inclusion of a more continuous measure of RSA in this study represents an important advancement over previous studies measuring infants’ physiological responses during the FFSF paradigm. Still, the current findings are limited in the same way as most previous FFSF findings in that they are based on a set of correlational associations. As such, causal inferences about the role of cardiac vagal tone or physiological synchrony in the development of emotion regulation remain premature. Currently, few studies in this literature are designed to address the causal relations between infants and mothers emotional states, but one important exception is the work by Waters and colleagues (Waters, West, & Mendes, 2014). In this study, mothers were randomly assigned to a stressful or non-stressful task before they were reunited with their 12- to 14-month-old infants. The results revealed that infants’ physiological reactivity mirrored the mothers’ physiological state and that this affect contagion was responsible for greater physiological synchrony between mothers and their infants throughout the remainder of the study.

One additional issue merits some consideration. Mothers in this study were instructed to not touch their infants. This prohibition may have decreased the magnitude of physiological synchrony between infants and mothers because touch is an important communication medium for mothers to share affective signals with their infants (e.g. Waters, West, Karnilowicz, & Mendes, 2017) and to facilitate emotion regulation (Lowe et al., 2016). Moreover, synchronous touching between infants and mothers during free play is associated with higher infant vagal tone (Feldman et al., 2010), underscoring how touch can influence autonomic activity. Future work should examine to what extent maternal touch interacts with physiological synchrony to influence infants’ biobehavioral regulation.

Another important area for future research is to investigate the contributions of physiological synchrony in more diverse samples varying along the dimensions of SES, race and ethnicity as well as risk status (e.g. Boeve, Beeghly, Stacks, Manning, & Thomason, 2019; Conradt & Ablow, 2010; Suurland et al., 2017). It is quite possible that physiological synchrony will play an even more important role in behavioral regulation for at-risk infants who may have regulatory difficulties.

## Conclusion

The implementation of the Still Face paradigm for studying infants’ emotion regulation has had a long and venerable history. Although infants typically display behavioral and physiological distress during the Still Face episode, which is followed by emotion regulation during the Reunion episode, there remain numerous confusions and contradictions in the literature (Moore & Calkins, 2004; Weinberg & Tronick, 1996). Based on the findings from the current study, it is apparent that individual differences in physiological reactivity and physiological synchrony between infants and mothers contribute to these inconsistent findings. Our results reveal that infants who are unable to suppress their vagal tone during the Still Face episode require greater co-regulation from their mothers to reduce their distress during the Reunion episode. Critically, this capacity for emotion regulation during Reunion is observed only for infants who were previously classified as engaging in positive physiological synchrony with their mothers during the Social Play episode. Infants classified as engaging in negative synchrony with their mothers were unable to regulate their negative affect during Reunion. By contrast, infants who were able to suppress their RSA and thus physiologically regulate their affective state during the Still Face were also able to regulate their emotional state during Reunion regardless of their physiological synchrony with their mothers during Social Play.

These findings suggest that physiological synchrony, like behavioral synchrony, is an informative index of the mother’s co-regulation of the infant’s affective state, but it seems to apply primarily to infants who are less capable of suppressing their vagal activity.

## Acknowledgments

We gratefully acknowledge all of the families who participated in this research. An outstanding team of students assisted with video coding and physiological data processing: Kelsey Blalock, Meghan Burmeister, Khushboo Chougule, Quinn Cox, Diandra Elsner, Deanne Hoaglund, Rebecca Hailperin-Lausch, Hannah Maluvac, Lela Minor, Pooja Pandita, Ellen Parrish, Marissa Radziwiecki, and Stephanie Younker. Karen Wilkie assisted with subject recruitment and testing. Dr. Stephen Porges and Dr. Gregory Lewis provided valuable feedback regarding the method and interpretation of the data.

## Footnotes

In previous iterations of this project, there was an attempt to create three synchrony categories (positive, negative, no synchrony) as a function of bootstrapped confidence intervals, but the outcome resulted in three unbalanced group sizes and reduced statistical power.

